# The microtubule minus end-binding protein CAMSAP2 does not regulate microtubule dynamics in primary pancreatic β-cells but facilitates insulin secretion

**DOI:** 10.1101/2022.05.05.490800

**Authors:** Kung-Hsien Ho, Anna B. Osipovich, Anissa Jayathilake, Mark A. Magnuson, Guoqiang Gu, Irina Kaverina

## Abstract

Glucose stimulation induces the remodeling of microtubules in Islet β-cells to potentiate glucose-stimulated insulin secretion. CAMSAP2 is a microtubule minus-end binding protein and is reported to stabilize and position microtubules in several non-β-cells, such as human retinal pigment epithelium cells. In immortalized insulinoma MIN6 cells, CAMSAP2 binds to and forms short stretches at microtubule minus ends in the cytoplasm, which is consistent with the reported subcellular localization and functions of CAMSAP2 in non-β-cells. Surprisingly, we found that CAMSAP2 expressed in primary islet β-cells does not form short stretches in the cytoplasm, but instead is localized to the Golgi apparatus. This novel localization is specific to β-but not α-cells in islets and it is independent of MT-binding. Knockdown of CAMSAP2 by shRNA impairs Golgi-ER trafficking, reduces total insulin content, and attenuates GSIS without affecting the MT dynamics or releasability of insulin granules in islet β-cells. Corresponding to these results, we found that primary islets and MIN6 cells express different CAMSAP2 isoforms. We propose that primary islet β cells use a novel CAMSAP2 isoform for a MT-independent non-canonical function, which is to promote Golgi-ER trafficking that supports efficient production of insulin secretory granules.

## Introduction

### 1. Insulin biosynthesis

Pancreatic islets are central regulators of glucose homeostasis. Increased blood glucose induces β-cells to secrete insulin in a process called glucose-stimulated insulin secretion (GSIS). The synthesis of insulin starts with its mRNA translated into preproinsulin (Chan et al., 1976). Preproinsulin is imported into the ER, where its signal peptide is rapidly cleaved to generate proinsulin. After proper folding and the formation of three intramolecular disulfide bonds, proinsulin is transported to the Golgi apparatus [referred as Golgi in short] (Orci, 1982). In the Golgi, proinsulin assembles into hexamers and is packed into secretory vesicles (Brill and Venable, 1968). Further proteolytic cleavage generates the C-peptide and mature insulin inside the vesicles (Hutton, 1982; Orci et al., 1986; Steiner et al., 1974).

### 2. GSIS and insulin pools

After a meal, high glucose transported into β-cells is metabolized to elevate intracellular ATP/ADP ratio, which induces the closure of ATP-sensitive potassium channels on the plasma membrane (Ashcroft et al., 1984; Cook and Hales, 1984; Dunne and Petersen, 1986; Kakei et al., 1986). This leads to increased trans-membrane potential and the opening of voltage-dependent calcium channel on plasma membrane (Dean and Matthews, 1970). The calcium influx facilitates the fusion of secretory vesicles to plasma membrane to release insulin into the blood stream (Bokvist et al., 1995). Islet β-cells store thousands of insulin vesicles in the cytoplasm. Traditionally, these vesicles are categorized into two separate pools, the readily releasable pool, RRP and the reserved pool, RP (Olofsson et al., 2002; Rorsman and Renstrom, 2003). Vesicles in the RRP are either docked to the plasma membrane or are in close vicinity (Ohara-Imaizumi et al., 2007; Shibasaki et al., 2007). They can be secreted immediately following the calcium influx and are largely released in the first phase of insulin secretion that lasts a few minutes (Curry et al., 1968; Ohara-Imaizumi et al., 2004). Vesicles in RP are localized away from the plasma membrane. They require long-ranged transportation to reach cell periphery and are responsible for the second phase of insulin secretion that can last for hours. Meanwhile, glucose metabolism also triggers new insulin granule biosynthesis that replenishes the diminishing insulin pools for sustainable β-cell function.

### 3. Microtubules in β-cells and polarity

The cytoskeletons, both actin filaments and microtubules, are key cellular structure that regulate insulin biosynthesis and secretion in β-cells. F-actin and myosin 5a promote the transportation of insulin vesicles from the cell interior to periphery (Ivarsson et al., 2005; Varadi et al., 2005). In the cell periphery, cortical actin forms a barrier underneath the plasma membrane. Glucose stimulation induces the reorganization of F-actin in cell periphery to facilitate insulin secretion (Thurmond et al., 2003; Wang and Thurmond, 2009). Microtubules comprise α- and β-tubulin heterodimers whose directional assembly confers the polarity of microtubules. The α-tubulin exposed end is called the minus end and the β-tubulin exposed end is the plus ends (Bergen and Borisy, 1980). The orientation of microtubules together with plus/minus end-directed motors determine the direction of vesicle transportation along microtubules (Vale and Milligan, 2000; Vale et al., 1985).

### 4. Microtubules regulate the transportation and secretion of insulin vesicles

Similar to the actin cytoskeleton, microtubules also have dual functions in the regulation of insulin secretion. Microtubule-dependent long-ranged transportation facilitates the delivery of insulin vesicles from the cell interior to close to the plasma membrane (Heaslip et al., 2014; Varadi et al., 2002). In the cell periphery, microtubules regulate the positioning of insulin vesicles through withdrawing them back to the cell interior, a process that counteracts the delivery of insulin vesicles from the cell interior to periphery (Zhu et al., 2015). When β-cells are excited with high glucose, enhanced microtubule remodeling, including increased depolymerization and nucleation (Heaslip et al., 2014; Zhu et al., 2015), increases the turnover leading to increased microtubule numbers and reduced length per microtubule (Muller et al., 2021). The remodeling of peripheral microtubules inhibits the net withdrawal and facilitates robust secretion by increasing the availability of insulin vesicles in cell periphery (Ho et al., 2020; Zhu et al., 2015).

### 5. CAMSAP2 is a MT minus end binding proteins, and can be used to visualize MT orientation in cells

CAMSAPs (<u>CA</u>l<u>M</u>odulin-regulated <u>S</u>pectrin-<u>A</u>ssociated <u>P</u>rotein) specifically bind to free microtubule minus ends (Goodwin and Vale, 2010; Meng et al., 2008). There are three homologues in mammals, CAMSAP1, 2, and 3 (Baines et al., 2009). In RPE1 (human retinal pigment epithelium) cells, CAMSAP2 stabilizes microtubule minus ends, while its growing recruits more CAMSAP2 to the nascent minus ends and overtime forming short stretches of CAMSAP2-decoration (Jiang et al., 2014). CAMSAP2 also facilitates the positioning of microtubule minus ends to the Golgi through interacting with Golgi-localized AKAP450 and myomegalin (Wu et al., 2016). The directionality of microtubules (the distribution of minus and plus ends) is important for directional long-ranged vesicle transportation. Understanding the directionality of microtubules in β-cells under different glucose concentration is important for constructing the dynamic model of microtubule-mediated insulin transportation and secretion. CAMSAP2 is a good marker to visualize the distribution of microtubule minus ends and could reveal the directionality of the microtubule network in β-cells. We planned to use this approach to label minus ends and follow their distribution responding to glucose stimulation. Surprisingly, we discovered that CAMSAP2 exhibit a novel subcellular localization to the Golgi in primary β-cells independent of binding to microtubules. Further analyses revealed CAMSAP2 is required for efficient Golgi-to-ER trafficking, which facilitates robust GSIS.

## Results

### CAMSAP2 is specifically enriched at the Golgi in islet β-cells

In several well characterized non-β-cells, such as Hela and RPE1, CAMSAP2 binds to the minus end of microtubules and form small stretches in the cytoplasm (Jiang et al., 2014). We used the immortalized mouse insulinoma MIN6 cells that possesses some β-cell secretary properties as a β-cell model and labeled CAMSAP2 by immunofluorescence staining as a surrogate to visualize the directionality of the microtubule network in β-cells (S1A-D). In interphase MIN6 cells, CAMSAP2 staining exhibits short stretches or puncta in the cytoplasm as in other previously reported non-β-cells. In dividing MIN6 cells at telophase, we observed CAMSAP2 signal at the distal tips of the mid-body, confirming the specificity of CAMSAP2 binding to microtubule minus ends in β-cells (S1E). A similar pattern of CAMSAP2 puncta/stretches in the cytoplasm was observed in the rat insulinoma INS-1 cells (S1F-H).

In primary β-cells inside mouse islets, however, we did not observe CAMSAP2-labeled short stretches in the cytoplasm, but instead, the CAMSAP2 signal is localized to the cell interior and overlaps with Golgi markers, such as GM130 and GCC185 (Fig. 1A-F). The other two homologues, CAMSAP1 and CAMSAP3, are not localized to the Golgi in primary islet β-cells (Fig. S1H-M). Therefore, we focus on CAMSAP2 and its functions in primary β-cells in this report.

**Figure 1.**
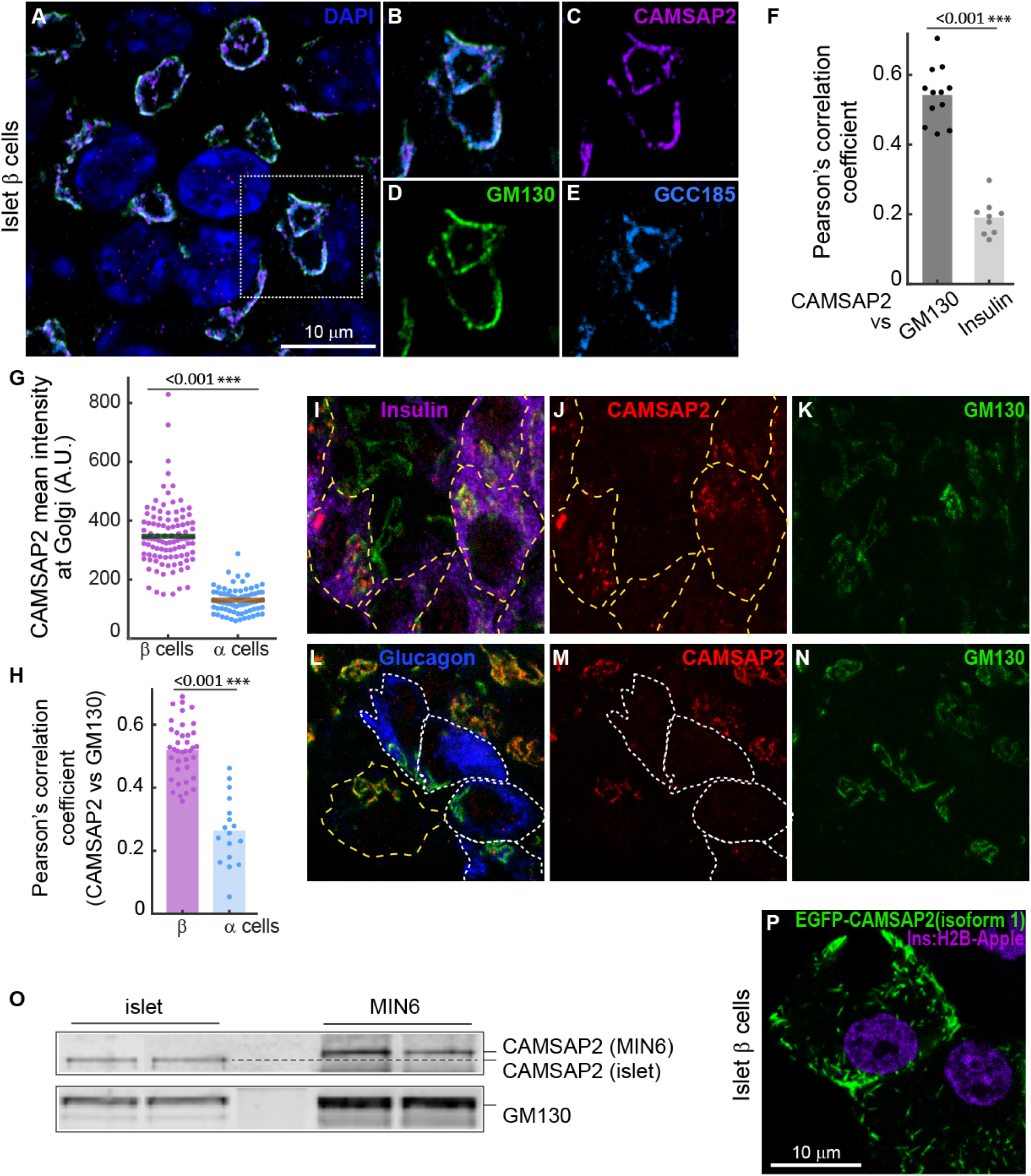
CAMSAP2 localizes to the Golgi in islet β-cells. (A-E) Immunofluorescence staining of GM130 (green), CAMSAP2 (magenta), and GCC185 (cyan) in β-cells in mouse islets. (F) Pearson’s correlation coefficient of CAMSAP2 vs GM130 and CAMSAP2 vs insulin immunofluorescence. Dots represent image sections of islets. Columns represent the average of three animals. *** p<0.01 (t test). (G) Quantification of CAMSAP2 immunofluorescence intensity at the Golgi in mouse islets. Dots represent the mean intensity of CAMSAP2 localized at the Golgi of individual cells. Bars represent averages from three animals. *** P<0.001 (t test). (H) Pearson’s correlation coefficient of CAMSAP2 vs GM130 immunofluorescence in mouse islets. Dots represent individual cells. Columns represent the average of three animals. *** p<0.01 (t test). (I-N) Immunofluorescence staining of insulin (magenta), glucagon (blue), CAMSAP2 (red), and GM130 (green) in mouse islets. (O) Immunoblotting against CAMSAP2 and GM130 in lysates from MIN6 and from islets. The dashed line indicates the position of CAMSAP2 from islet lysates. (P) Fluorescence images of ectopically expressed EGFP-CAMSAP2 (human CAMSAP2 isoform 1, green) in mouse islets. Red, *Ins2* promoter-driven H2B-mApple.

Immortal β-cells like MIN6 and INS-1 do not have the cell-cell interactions with other type of islet endocrine cells when compared to primary β-cells inside islets, which has been shown to influence cytoskeletal configuration (Liu et al., 2014). We examined the subcellular localization of CAMSAP2 in primary β-cells dissociated from isolated mouse islets and observed enriched CAMSAP2 at the Golgi similar to that in β-cells inside islets (Fig. S1N-P). These results suggest that the unique localization of CAMSAP2 is intrinsic to primary β-cells and is not due to external influence from other cells in islets.

We next asked whether another important islet endocrine cells, α-cells, also have CAMSAP2 enriched at the Golgi. We labeled β- and α-cells in islets with immunofluorescence staining against insulin and glucagon respectively and compared the intensity of CAMSAP2 at the Golgi. We found that β-cells have significantly higher CAMSAP2 level at the Golgi than α-cells (1G-N). These results suggest that the localization of CAMSAP2 at the Golgi is not a general feature in primary islet endocrine cells. We hypothesize that this unique localization of CAMSAP2 may be associated with some β-cells specific functions and tested it in following paragraphs. Interestingly, when compared to MIN6 cells, CAMSAP2 expressed in islet β-cells exhibits a faster electrophoretic mobility with an estimated molecular weight difference of around 10.6 KDa (Fig. 1O). These results suggest that primary β-cells express a different CAMSAP2 variant, which may confer its unique localization to the Golgi. To test this hypothesis, we ectopically expressed an EGFP-tagged human CAMSAP2 isoform 1, which is the longest reported variant, in mouse islet β-cells. We found that the EGFP signal forms small stretches in the cytoplasm, which is similar to the endogenous CAMSAP2 expressed in MIN6 cells (Fig. 1P, S1A-E). These results suggest that the long CAMSAP2 isoform binds to microtubule minus ends in islet β-cells and support the hypothesis that they express a different CAMSAP2 variant.

### The CAMSAP2 signal detected at the Golgi in islet β-cells is specific

To further confirm the detection of CAMSAP2 using the antibodies is specific, we ectopically expressed EGFP-tagged human CAMSAP2 isoform 1 in MIN6 and compared it to the signal detected by anti-CAMSAP2 immunofluorescence. We observed strong co-localization of the EGFP and the CAMSAP2 immunofluorescence signal (Fig. 2A). We also tested the specificity of the anti-CAMSAP2 antibody in MIN6 cells expressing EGFP-tagged CAMSAP1 or CAMSAP3. We found that the EGFP signal from either CAMSAP1 or CAMSAP3 do not overlap with the CASAMP2 immunofluorescence signal (Fig. 2B-C). To further verify the specificity of the antibody detection of CAMSAP2 in islet β-cells, we used shRNA to deplete CAMSAP2. Compared to the non-targeting shRNA, islet cells expressing CAMSAP2-targeting shRNA showed reduced CAMSAP2 immunofluorescence signal (Fig. 2D-L). These results suggest that the anti-CAMSAP2 antibodies used in this study is specific in detecting CAMSAP2 in islet β-cells.

**Figure 2.**
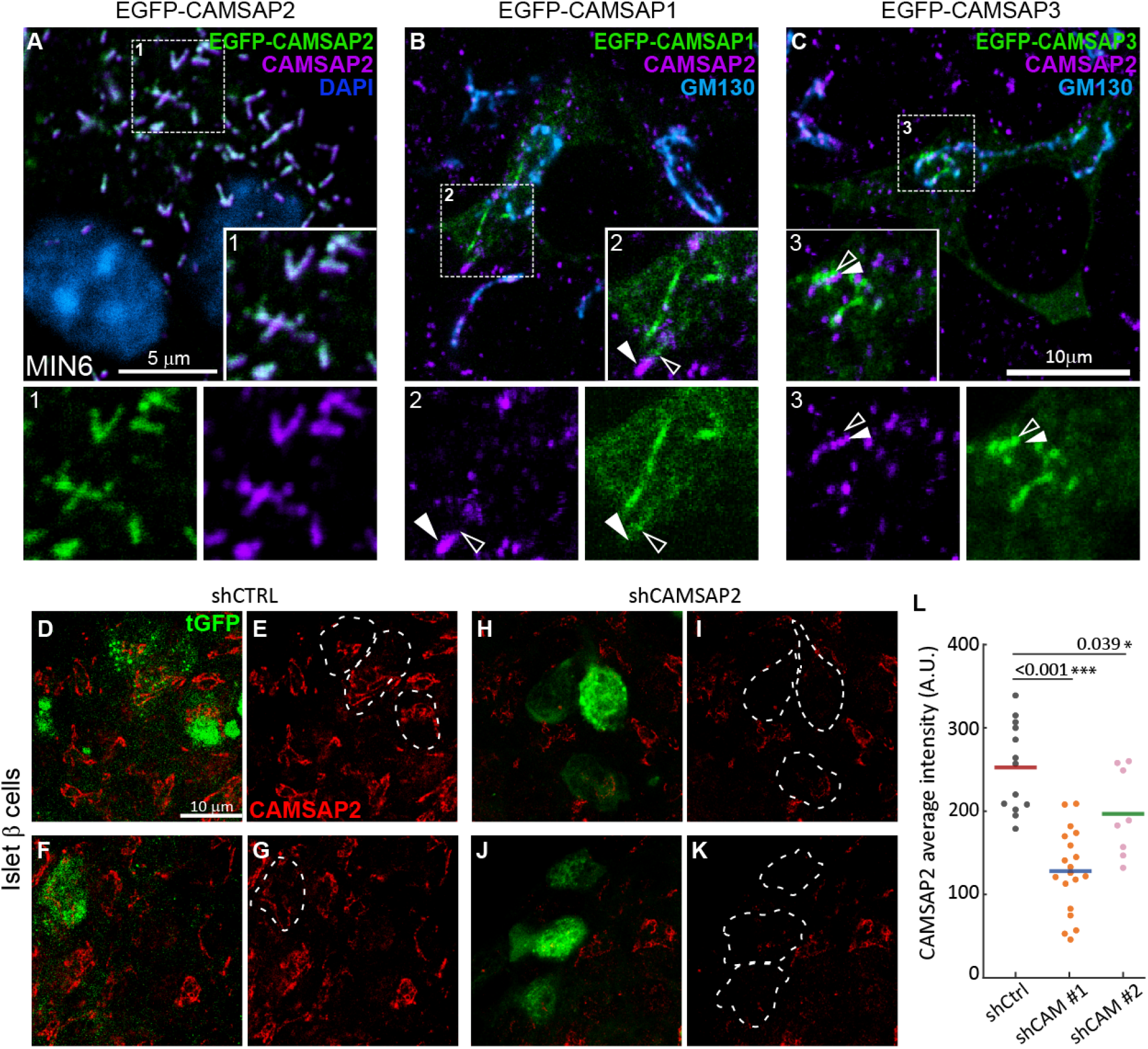
The Immunofluorescence staining detects CAMSAP2 specifically and islet β-cells express a smaller CAMSAP2 isoform. (A) Immunofluorescence staining of CAMSAP2 (magenta) in MIN6 ectopically expressed EGFP-CAMSAP2 (human isoform 1, green). Blue, DAPI. (B-C) Immunofluorescence staining of CAMSAP2 (red) and GM130 (magenta) in MIN6 ectopically expressed EGFP-CAMSAP1 (B) or EGFP-CAMSAP3 (C) in green. Closed arrowheads, CAMSAP2; open arrowhead, EGFP-CAMSAP1 or EGFP-CAMSAP3. (D-K) Immunofluorescence staining of CAMSAP2 (red) in islet cells expressing non-targeting shRNA (shCtrl, D-G) or shRNA targeting CAMSAP2 #1 (shCAMSAP2, H-K). Green, tGFP to indicate successful transduction. (L) Quantification of CAMSAP2 intensity in transduced β-cells in mouse islets. * p<0.05, *** p<0.001 (Dunnett’s multiple comparisons test).

### The localization of CAMSAP2 to the Golgi is independent of binding to microtubules

The Golgi apparatus is the major microtubule nucleation site in islet β-cells (Zhu et al., 2015). CAMSAP2 is reported to anchor microtubule minus ends to the Golgi through its interactions with AKAP450 and myomegalin (Wu et al., 2016). It is possible that the Golgi-localized CAMSAP2 in islet β-cells may represent a highly organized and concentrated minus ends anchored to the Golgi. To test this hypothesis, we used nocodazole to disassemble microtubules and then removed it to allow the re-growth of microtubules from the Golgi. After the removal of nocodazole, the abundance of CAMSAP2 at the Golgi exceeds the number of microtubules emerged from the Golgi and thus the number of minus ends (Fig. 3A-D). Nocodazole treatment induces microtubules depolymerization and the disassembly of the Golgi apparatus into smaller mini-stacks. We found that the fold enrichment of CAMSAP2 at the Golgi over cytoplasm is not altered in nocodazole-treated cells when compared to that in untreated ones (Fig. 3E-L & S). These results suggest that microtubule depolymerization does not affect the localization of CAMSAP2 to the Golgi.

**Figure 3.**
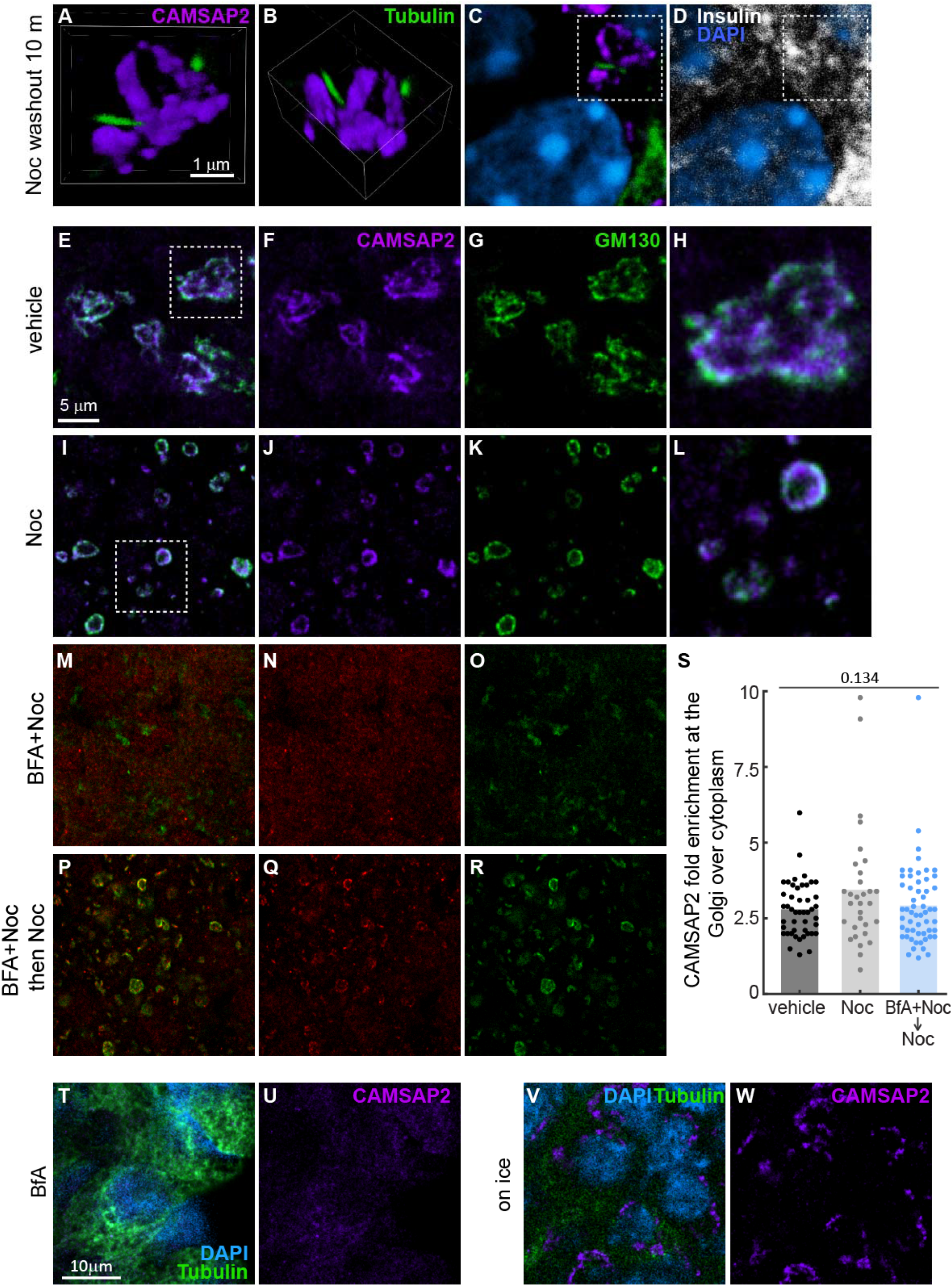
The localization of CAMSAP2 to the Golgi in islet β-cells is independent of binding to microtubules. (A-D) Immunofluorescence staining of CAMSAP2 (magenta), Tubulin (green), and insulin (white) in mouse islets. Islets were incubated with nocodazole to depolymerize microtubules and the nocodazole was removed to allow microtubule growth. (A-B) 3D reconstituted images of Golgi mini-stacks. Blue, DAPI. (E-R) Immunofluorescence staining of GM130 (green) and CAMSAP2 (magenta) in mouse islets treated with nocodazole (I-L), nocodazole plus Brefeldin A (M-O), and washed and incubated with nocodazole without Brefeldin A (P-R). (S) Quantification of CAMSAP2 fluorescence intensity fold enrichment at the Golgi over that in the cytosol. Dots represent individual cells in mouse islets. Columns represent average of three animals (One-way ANOVA). (T-W) Immunofluorescence staining of tubulin (green) and CAMSAP2 (magenta) in mouse islets treated with Brefeldin A (R-S) or incubated on ice (T-U). Blue, DAPI.

It is possible that CAMSAP2 binds to and stabilizes small microtubule fragments at the Golgi and protects them from the nocodazole treatment. To further test whether binding to microtubules is required for the localization of CAMSAP2 to the Golgi, we used Brefeldin A, which reversibly blocks the endoplasmic reticulum (ER)-to-Golgi membrane trafficking resulting in the collapse of the Golgi (Niu et al, 2005). After the Brefeldin A treatment, CAMSAP2 is no longer localized to the cell interior, which is consistent with the findings that CAMSAP2 localizes to the Golgi (Fig. 3M-O). The blocking of ER-to-Golgi trafficking can be lifted by removing Brefeldin A, which allows the Golgi to re-form in the cytoplasm (Fig. 3M-R). When nocodazole is present in both the treatment and the removal of Brefeldin A, we still observe the localization of CAMSAP2 to nascent Golgi mini-stacks at a level of enrichment comparable to that of untreated islet β-cells (Fig. 3S). The total protein level of CAMSAP2 is not altered by different concentration of glucose, or the treatment of nocodazole or Brefeldin A (Fig. S2). These results suggest that the localization of CAMSAP2 at the Golgi is not dependent of microtubule-binding.

Although the Brefeldin A treatment abolishes the localization of CAMSAP2 in the cell interior, it does not induce microtubule disassembly (Fig. 3T-U). These results suggest that the Brefeldin A treatment induced loss of CAMSAP2 signal in the cell interior is not due to disruption of the microtubule network. Additionally, cold treatment induces microtubule disassembly but does not immediately disrupt the Golgi structure because of the slow-down of membrane trafficking at low temperature. In this condition, CAMSAP2 signal remains enriched in the cell interior in the absence of microtubules (Fig. 3V-W). Together, these results indicate that the localization of CAMSAP2 to the Golgi is independent of microtubule-binding.

### Depletion of CAMSAP2 in islet β-cells impairs insulin secretion

The findings that the Golgi-localization of CAMSAP2 is specific to islet β-cells but not to α-cells suggest that CAMSAP2 may be required for the functions of β-cells. To test this hypothesis, we used lentivirus to deliver shRNAs to islet cells for specific depletion of CAMSAP2. To reach a high transduction rate, we dissociated cells from isolated mouse islets, mixed them with the lentivirus and allowed cells to re-aggregate into pseudoislets (Fig. 4A-B). The cell-cell interactions inside pseudoislets create a niche similar to inside an intact islet and is a good platform to access the secretion capacity of β-cells (Reissaus and Piston, 2017). The expression of either of the two CAMSAP2-targeting shRNAs does not alter the basal insulin secretion in low glucose conditions, but impairs GSIS (Fig. 4C). Liraglutide, a GLP-1 analogue, enhances GSIS of CAMSAP2-depleted pseudoislets, but it is still significantly lower than its control counterpart. These results suggest that CAMSAP2 is required for robust secretion induced by high glucose.

**Figure 4.**
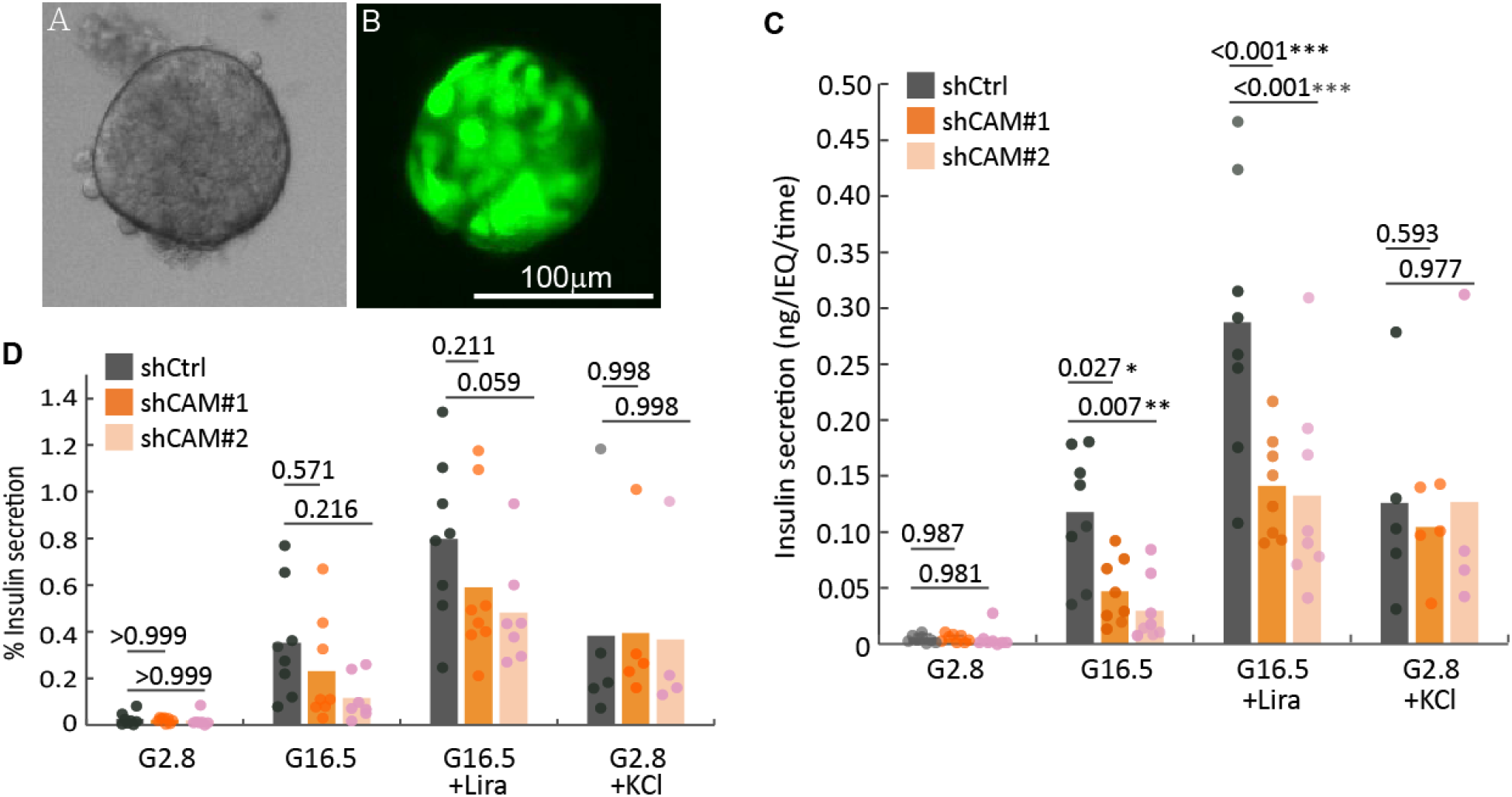
CAMSAP2 knockdown in islet β-cells impairs GSIS but not KSIS. (A-B) Representative widefield (A) and fluorescence (B) images of a pseudoislet generated from dissociated islet cells transduced with lentivirus. The lentivirus contain shRNA and a tGFP expression cassette. (C) GSIS and KSIS of pseudoislets expressing non-targeting (shCtrl) or CAMSAP2-targeting (shCAM) shRNA. * p<0.05, ** p<0.01, *** p<0.001 (Holm-Šídák’s multiple comparisons test). Pseudoislets were sequentially incubated with 2.8 or 16.5 mM glucose for 45 minutes and in 2.8 mM glucose plus 25 mM KCl for 30 minutes. Secretion were normalized by total IEQ. (D) Percent secretion of shRNA expressing pseudoislets calculated as a percentage of total insulin content (Holm-Šídák’s multiple comparisons test).

A high level of extracellular KCl induces membrane depolarization and the release of insulin vesicles that are either docked or close to the plasma membrane (Ohara-Imaizumi et al., 2002). When stimulated with KCl, the secretion of pseudoislets expressing CAMSAP2-targeting shRNAs is comparable to ones expressing the non-targeting shRNA (Fig. 4C). These results suggest that the reduced GSIS in CAMSAP2 depletion cells is likely due to impaired vesicle transportation. Consistently, when we compare GSIS as a percentage of total insulin content between pseudoislets expressing CAMSAP2-targeting and control shRNAs, the difference in secretion is reduced and is not statistically significant (Figure 4D). These results suggest that the capacity of CAMSAP2-depleted pseudoislets to secrete insulin in response to glucose stimulation is not significantly weaker than the controls. We hypothesize that the reduction in the absolute amount of secreted insulin in CAMSAP2-depleted pseudoislets is likely due to a changed total insulin content and tested it below.

### Depletion of CAMSAP2 in islet β-cells leads to reduced total insulin and partially impairs the ER-to-Golgi membrane trafficking

To understand how CAMSAP2 depletion influences insulin secretion in β-cells, we first tested whether the synthesis of insulin is affected. We examined the steady-state level of proinsulin by immunofluorescence staining and found that CAMSAP2 depletion does not change the level of proinsulin (Fig. S3), suggesting that the translation and folding of proinsulin are not changed. We next examined the level of total insulin in CAMSAP2-depleted islet β-cells. We found that the majority of islet β-cells transduced by the CAMSAP2-targeting lentivirus showed a moderate reduction in total insulin level. Although we observed a few transduced cells that have strong insulin staining, the average of total insulin content from cells expressing CAMSAP2-targeting shRNA is significant lower than that of control cells expressing non-targeting shRNA (Fig. 5A-H).

**Figure 5.**
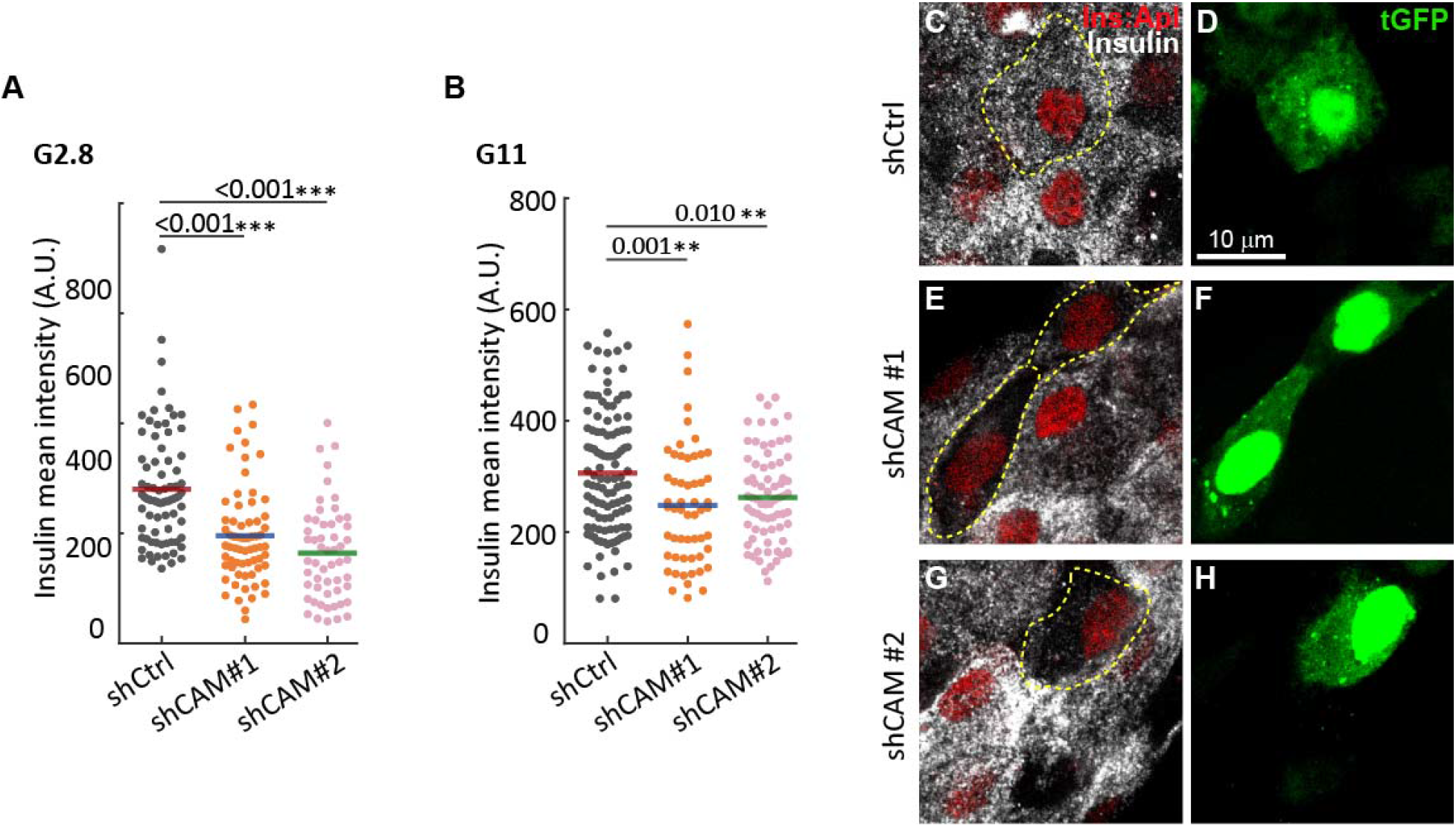
CAMSAP2 depletion reduces total insulin content in islet β-cells (A-B) Quantification of immunofluorescence intensity of insulin in β-cells in mouse islets cultured in media containing 2.8 mM (A) or 11 mM (B) glucose. Dots represent the mean intensity of total insulin of individual cells. Bars represent averages from three animals. ** p<0.01, *** p<0.001 (Dunnett’s multiple comparisons test). (C-H) Immunofluorescence staining of insulin (white) in islets transduced with shRNAs and cultured in media containing 11 mM glucose. Dashed lines delineate transduced cells. Red, *Ins2* promoter-driven H2B-mApple; Green, tGFP to indicate successful transduction.

Proinsulin is transported from the ER to the Golgi for further assembly into hexamers and package into insulin vesicles. Because CAMSAP2 is enriched at the Golgi in primary islet β-cells, we tested the hypothesis that CAMSAP2 depletion may impact the trafficking between the Golgi and ER by using Brefeldin A to challenge the ER-to-Golgi membrane trafficking. Brefeldin A inhibits the ER-to-Golgi trafficking (anterograde), and the retrograde transportation/direct fusion results in the reduction of the Golgi overtime. The *cis*-Golgi matrix protein, GM130, is not transported to the ER after Brefeldin A treatment and is released into the cytoplasm as the Golgi collapse (Nakamura et al, 1995). We monitored the level of GM130 in the cytoplasm as a readout of the speed of the Golgi-to-ER retrograde transportation. We found that CAMSAP2 depletion significantly slows the increase of GM130 level in the cytoplasm when compared to the control cells expressing non-targeting shRNA (Fig. 6A-Q). These results suggest that CAMSAP2-depletion partially impairs the Golgi-to-ER trafficking in islet β-cells. When Brefeldin A is removed, the reappearance and expansion of the Golgi overtime leads to a gradual reduction of GM130 level in the cytoplasm, which is used as a readout for the ER-to-Golgi transportation. We found that CAMSAP2-depleted cells have a slower reduction of cytoplasmic GM130 level after 90 minutes of Brefeldin A removal compared to the controls (Fig. 6R-V). Together, these results suggest that CAMSAP2 depletion impairs the trafficking between the Golgi and ER, which likely contributes to the reduced insulin content. We speculate that this may impact the transportation of insulin vesicles and the replenishment of the RRP in islet β-cells, leading to weaker GSIS.

**Figure 6.**
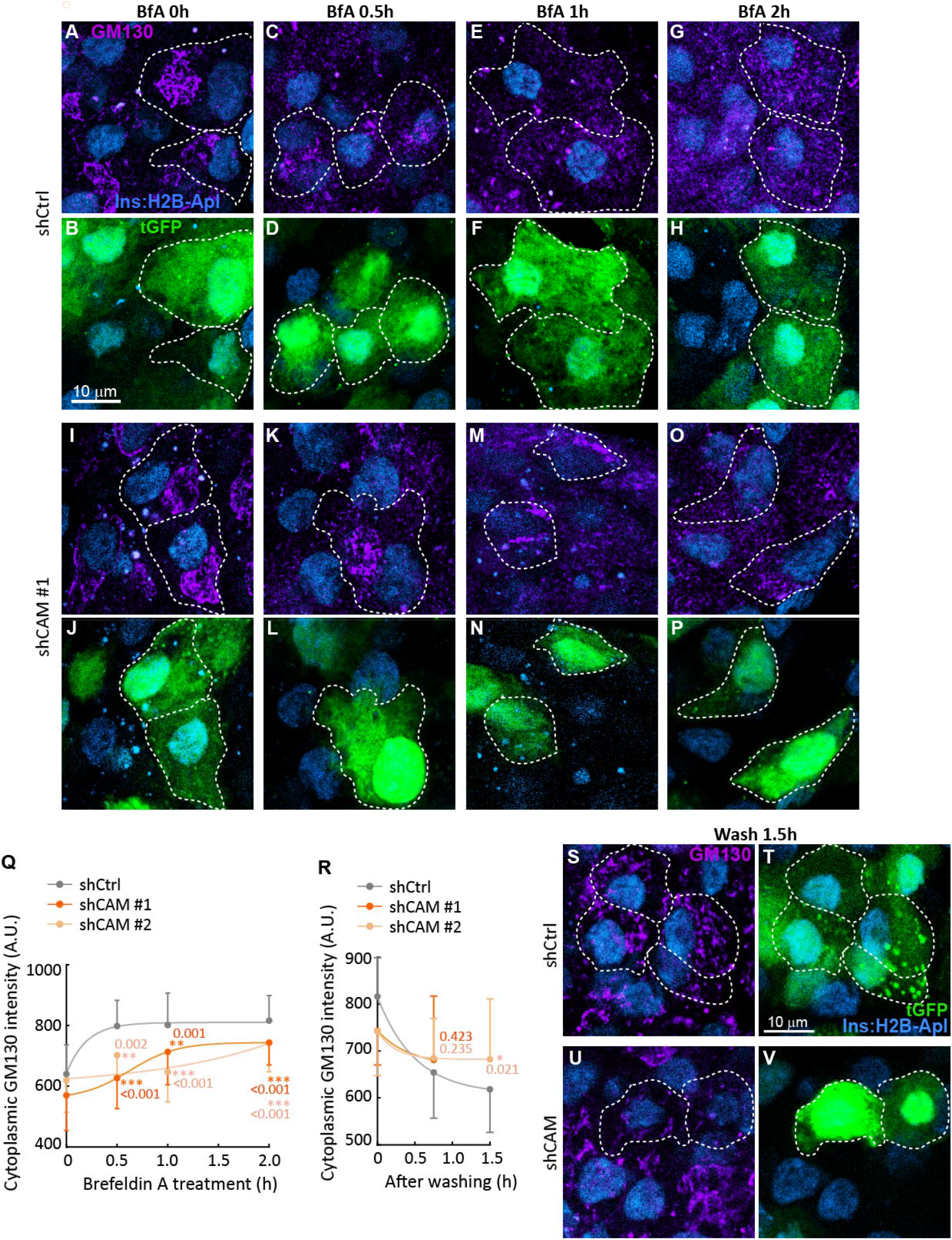
CAMSAP2 depletion impairs the ER-Golgi trafficking in islet β-cells (A-P) Immunofluorescence staining of GM130 (magenta) in islets transduced with shRNAs treated with Brefeldin A. Dashed lines delineate transduced cells. Cyan, *Ins2* promoter-driven H2B-mApple; Green, tGFP to indicate successful transduction. (Q-R) Quantification of cytoplasmic GM130 immunofluorescence intensity of β-cells in mouse islets in Brefeldin A treatment (Q) and after Brefeldin A removal (R). The cytoplasmic GM130 intensity is measured to exclude the Golgi and normalized by the mean GM130 intensity of the entire cell. Dots represent the average from three animals. Bars represent standard deviation. ** p<0.01, *** p<0.001 (Dunnett’s multiple comparisons test). (S-V) Immunofluorescence staining of GM130 (magenta) in islets transduced with shRNAs, treated with Brefeldin A, and incubated in fresh media to remove Brefeldin A. Dashed lines delineate transduced cells. Cyan, *Ins2* promoter-driven H2B-mApple; Green, tGFP to indicate successful transduction.

### CAMSAP2 does not influence glucose-induced microtubule dynamics in islet β-cells

The coupling of microtubule dynamics to glucose stimulation in islet β-cells is an important mechanism to fine-tune the dosing of insulin secretion. High glucose stimulates the disassembly of microtubules in islet β-cells to facilitate robust secretion. CAMSAP2 is known to binds to and stabilizes microtubule minus ends. It is possible that the depletion of CAMSAP2 in islet β-cells may also disrupt the coupling of microtubule disassembly to high glucose stimulation contributing to impaired GSIS. To test this hypothesis, we examine the level of detyrosinated tubulin in islet cells expressing CAMSAP2-targeting shRNA. The C-terminal last tyrosine of α-tubulin in microtubules is removed by tubulin carboxypeptidase generating detyrosinated tubulin, whose abundance is a well-established indicator of long-lived microtubules (Gundersen et al., 1987). We found that the depletion of CAMSAP2 does not influence the reduction of detyrosinated tubulin following glucose stimulation and there is no difference when comparing the level of detyrosinated tubulin between control and CAMSAP2-depleted β-cells incubated with either 2.8 or 11 mM glucose (Fig. S4A-M). These results suggest that CAMSAP2 is not required for the glucose-induced microtubule disassembly. To further test whether CAMSAP2 depletion affects the microtubule network in islet β-cells, we used methanol extraction to remove cytosolic free tubulin and immunostained for microtubules. We did not observe a significant difference in the abundance of microtubules between islet β-cells expressing CAMSAP2-targeting or the control shRNAs (Fig. S4N-S). The distribution of microtubules in the cytoplasm is also not altered by CAMSAP2 depletion (Fig. S4T). To conclude, CAMSAP2 depletion does not change the microtubule dynamics responding to glucose stimulation or the overall abundance of microtubules in islet β-cells.

## Discussion

### 1. Primary β-cells and immortalized insulinoma cells express different variants of CAMSAP2 with different subcellular localization and functions

Immortalized insulinoma cells, such as MIN6 and INS-1, are commonly used models to study β-cell biology. In both cell lines, CAMSAP2 binds to microtubule minus-ends and forms short stretches in the cytoplasm. We report here that primary β-cells express a different variant of CAMSAP2, which has unique subcellular localization and functions. Instead of forming short stretches in the cytoplasm, CAMSAP2 expressed in primary β-cells is specifically enriched at the Golgi independent of binding to microtubules. In other cell types, such as RPE1 and human mammary epithelial cells (hTMECs), CAMSAP2 stretches are reported to predominantly localize close to the Golgi (Wu et al., 2016). These represent CAMSAP2-bound microtubule minus ends that are organized at the Golgi. In contrast, the Golgi-localized CAMSAP2 in primary islet β cells does not form short stretches and is independent of binding to microtubules. This unique localization is likely important to the moonlight functions of CAMSAP2 in primary β-cells to support Golgi-to-ER trafficking and likely facilitates the generation of insulin vesicles. When CAMSAP2 is depleted, β-cells have slower Golgi-to-ER trafficking, reduced total insulin content, and partially impaired GSIS. Our findings of the differential variant and functions of CAMSAP2 between insulinoma cells and primary β-cells suggest that researchers should be mindful when using immortalized insulinoma cells as a model to study particular proteins and cell biology of primary β-cells. It is possible that they may express a different variants of the protein of interest with different cellular functions when compared primary β-cells.

### 2. CAMSAP2 was identified in as a type 1 diabetes susceptibility locus in GWAS

CAMSAP2 was originally identified as an interaction partner of calmodulin and spectrin and was later discovered to be a microtubule minus end-binding protein. Interestingly, GWAS has mapped a type 1 diabetes susceptibility locus to CAMSAP2 at an intron, but the biological connection between CAMSAP2 and T1D was unknown (Onengut-Gumuscu et al., 2015). Our findings that CAMSAP2 is required for efficient insulin production and GSIS provide a possible explanation to its association with T1D. It is possible that the identified mutation of CAMSAP2 from GWAS might influence Golgi membrane trafficking and/or insulin vesicle generation leading to increased susceptibility to type 1 diabetes. By contrast, the other two homologues CAMSAP1 and CAMSAP3, which do not localize to the Golgi in primary β-cells, were not linked to diabetes in GWAS.

### 3. CAMSAP2 and MT

In primary β-cells, we did not observe CAMSAP2 forming cytoplasmic stretches and we did not detect the slightly longer CAMSAP2 variant expressed in MIN6 cells from immunoblotting. Although, we cannot exclude the possibility that primary β-cells may express the long variant found in MIN6 at a lower level and binds to microtubule minus ends, it is likely that the other two homologues, CAMSAP1 and CAMSAP3 are the main microtubule minus end-binding protein in primary β-cells.

When we ectopically expressed the human CAMSAP2 isoform 1 (the longest reported variant), we detected short stretches in the cytoplasm and not in the Golgi. These short stretches are likely CAMSAP2-decorated microtubule minus ends. Interestingly a large portion of those CAMSAP2 stretches are in cell periphery. This distribution of microtubule minus ends is different from other non-β-cells, such as RPE1, which has CAMSAP2 stretches predominantly localized close to the Golgi (Wu et al., 2016). When binds to minus ends, CAMSAP2 is reported to stabilize microtubules. We found that depletion of CAMSAP2 does not alter microtubule dynamics in respond to glucose stimulation in islet β-cells. These findings in primary β-cells are consistent with the model that the Golgi localized endogenous CAMSAP2 variant is not involved in binding to microtubule minus ends or tethering them to the Golgi.

### 4. Potential mechanism of how CAMSAP2 supports robust insulin secretion

We report here that CAMSAP2 is required for robust GSIS but not for KCl-induced secretion. These results suggest that CAMSAP2 is no required for the electrogenic mechanism of β-cells and the release of insulin that are already docked or close to the plasma membrane. Depletion of CAMSAP2 in β-cells leads to reduced total insulin content, which likely impacts the replenishing of insulin and the second phase of GSIS. We found that CAMSAP2 is specifically enriched at the Golgi in islet β-cells and its depletion impairs Golgi-to-ER trafficking. Rab GTPases and their regulatory guanine nucleotide exchange factors (GEF) are essential for the targeting of vesicles in Golgi-to-ER trafficking (Brandizzi and Barlowe, 2013). Two large scale studies using affinity purification and proximity-dependent biotinylation respectively have identified several Rab GEF as CAMSAP2-interaction partners, including DENND1A, DENND4C (Boldt et al., 2016), and DENND6A (Go et al., 2021). We speculate that CAMSAP2 may facilitate the localization and/or functions of these GEFs at the Golgi to promote efficient Golgi-to-ER trafficking in primary β-cells.

Together, we proposed that CAMSAP2 is required for efficient insulin vesicles generation and secretion. Its depletion impairs the trafficking between the Golgi and ER, reduces the total insulin content, which may slow the replenishment of insulin vesicles to the RRP. As a results, CAMSAP2 depleted cells showed reduced secretion induced by high glucose. In conclusion, our findings that CAMSAP2 is specifically enriched at the Golgi in islet β-cells and its depletion impairs GSIS provide a molecular connection of CAMSAP2 to diabetes.

## Materials and Methods

### 1. Mice

Mice were fed a regular chow diet *ad libitum* and maintained on a 12-12h dark-light cycle. Isoflurane inhalation was used for euthanasia. Wild-type CD-1 (ICR) mice were from Charles River Laboratories (Wilmington, MA). *Ins2^Apple^* mice have previously been described (Stancill et al., 2019). All Mouse experimentation followed protocols approved by the Vanderbilt University Institutional Animal Care and Use Committee.

### 2. Islet isolation and islet/cell culture

Islets were isolated from 8- to 12-week-old mice followed the published procedure (Li et al., 2009). Briefly, pancreas was perfused through the common bile duct with 2 mL of 0.8 mg/mL collagenase P (Roche, Indianapolis, IN) in Hanks’ balanced salt solution [HBSS] with Ca^2+^ and Mg^2+^ (Corning, Corning, NY). The dilated pancreas was digested at 37°C for 20 min. The homogenate was washed HBSS four times and islets were handpicked into islet media-G11 {RPMI-1640 media with 11 mM glucose (Gibco, Dublin, Ireland) plus 10% heat-inactivated Fetal Bovine serum [HI-FBS] (Atlanta Biologicals, Flowery Branch, GA)} and cultured at 37°C with 5% CO2. Primary β cells were harvested from dissociation of isolated islets (see below in the pseudoislet generation procedure) and culture in islet media-G11 in a glass-bottom 35-mm dish (MatTek, Ashland, MA) coated with human extracellular matrix [ECM] (Corning). Primary β cells were cultured over one night before fixation for imaging. MIN6 cells were grown in DMEM with 25 mM glucose (Gibco), 0.071 mM β-mercaptoethanol (Merck, Kenilworth, NJ), 10% HI-FBS, and 100 U/mL penicillin-100 μg/mL streptomycin (Gibco).

### 3. Small molecules, chemicals, and treatment

Liraglutide was from MedChemExpress (Monmouth Junction, NJ). Nocodazole, Brefeldin A, DMSO, KCl and other chemicals were from Merck. For the nocodazole treatment, isolated islets were incubated in islet media-G11 plus 10 mg/mL Nocodazole for 2 hours. For the Brefeldin A treatment, isolated islets were incubated in islet media-G11 plus 10 μg/mL Brefeldin A for 2 hours, transferred to islet media-G11 plus 10 mg/mL Nocodazole three times and incubated for two hours.

### 4. shRNA sequence

The CAMSAP2-targeting shRNA [shCAM] no #1 [TRCN0000252418, 5’-TTGTCCGGCTAGAGGATATTT-3’] and no #2 [TRCN0000252419, 5’-AGTTTCTCTGTCCGATTTAAA-3’] are in the plasmid backbone pLKO.1-CMV-tGFP and were from Sigma Millipore (St Louis, MO). The non-targeting shRNA control [shCtrl] plasmid, pLKO.1-shCtrl-puro, was from Addgene (cat. # 1864).

### 5. Islet dissociation, pseudoislets generation and transduction

To dissociate islets to primary cells, isolated mouse islets were recovered in islet media-G11 overnight, washed with HBSS without Ca2+ and Mg2+ for three times, digested with Accumax (Innovative Cell Technologies, San Diego, CA) at 30oC for 10 minutes, and gently pipetted up and down for 10 times. The cell suspension was spun at 1000g for 3 minutes and re-suspended in islet media-G11 mixed with concentrated lentivirus and DEAE-Dextran hydrochloride (Sigma Millipore). The mixture was loaded to the GravityPLUS hanging drop 96-well microplate (InSphero, Schlieren, Switzerland) and incubated for 4 days to generate pseudoislets. Pseudoislets were transferred to an ECM-coated 96-well plate and cultured for 1 day before analyses.

### 6. GSIS assay

The GSIS assay follows the published procedure (14). Briefly, pseudoislets were washed and conditioned in KRB buffer (111 mM NaCl, 4.8 mM KCl, 1.2 mM MgSO_4_, and 1.2 mM KH_2_PO_4_, 2.3 mM CaCl_2_, 25 mM NaHCO_3_, 10 mM HEPES and 0.2% BSA) plus 2.8 mM glucose (KRB-G2.8) for 1 hour, and then incubated with KRB-G2.8 for 1 hour to assess basal secretion. KRB buffer plus 16.5 mM glucose (KRB-G16.5) was used to incubate for 45 minutes to assess glucose-stimulated secretion. Pseudoislets were then incubated with KRB-G16.5 and 20 μM Liraglutide for 45 minutes. Pseudoislets were washed and incubated with KRB-G2.8 for 45 minutes and then stimulated with KRB-G2.8 and 30 mM KCl for 30 minutes. Secreted insulin was quantified using ELISA kits from ALPCO (Salem, NH).

### 6. Immunofluorescence staining and microscopy

For CAMSAP2 detection, samples were fixed with methanol at −20°C for 10 min. For the co-staining of CAMSAP2 with insulin or glucagon, samples were briefly fixed with 4% paraformaldehyde for 5 minutes and then fixed with methanol at −20°C for 10 min. For all other antigens, samples were fixed with 4% paraformaldehyde plus 0.1% saponin (MilliporeSigma, Burlington, MA). Antibodies used are as follows: anti–a-tubulin (cat. #ab18251, Abcam, Cambridge, U.K.), anti-E-cadherin (#610181, BD Biosciences, San Jose, CA), CAMSAP1 (#NBP1-26645, Novus Biologicals, Centennial, CO), CAMSAP2 (#NBP1-21402, Novus Biologicals), CAMSAP3 (#PA5-48993, Thermo Fisher Scientific, Waltham, MA), anti–detyrosinated tubulin (#AB3201, MilliporeSigma), anti-glucagon (#G2654, MilliporeSigma), anti-GM130 (#610823, BD Biosciences), antiinsulin (#A0564, Dako, Santa Clara, CA), anti-proinsulin (#GS-9A8-C, Developmental Studies Hybridoma Bank, Iowa City, IA), Secondary antibodies are from Invitrogen (Grand Island, NY). Vectashield Mounting Medium was from Vector Labs (Burlingame, CA). Confocal Images were captured using Nikon Eclipse A1R laser scanning confocal microscope equipped with a CFI Apochromat TIRF 1003/1.45 oil objective. Widefield images were captured using EVOS FL microscope equipped with an LPlan PH2 203/0.4 objective. Images were quantified using Nikon NIS-Elements. Mean intensity (average fluorescence intensity per pixel after background subtraction) of the epitope was measured. Background subtraction used the average intensity of a region without cells.

### 7. Immunoblotting

Isolated mouse islets and MIN6 cells were lysed in Tris-lysis buffer (10 mM Tris-Cl pH 7.5, 100 mM NaCl, 1% Triton X-100, 10% glycerol, and cOmplete™ protease inhibitor cocktail [Roche, Basel, Switzerland]) on ice. 75 μg of total protein in the lysate was loaded per lane and it was resolved in a 10% acrylamide gel. Antibodies used are as follows: IRDye 800 goat anti-rabbit IgG (Rockland Immunochemicals, Pottstown, PA) and IRDye 700DX goat anti-mouse IgG (LI-COR, Lincoln, NE). Blots were imaged using the Odyssey CLx imager (LI-COR).

---

**Figure S1.**
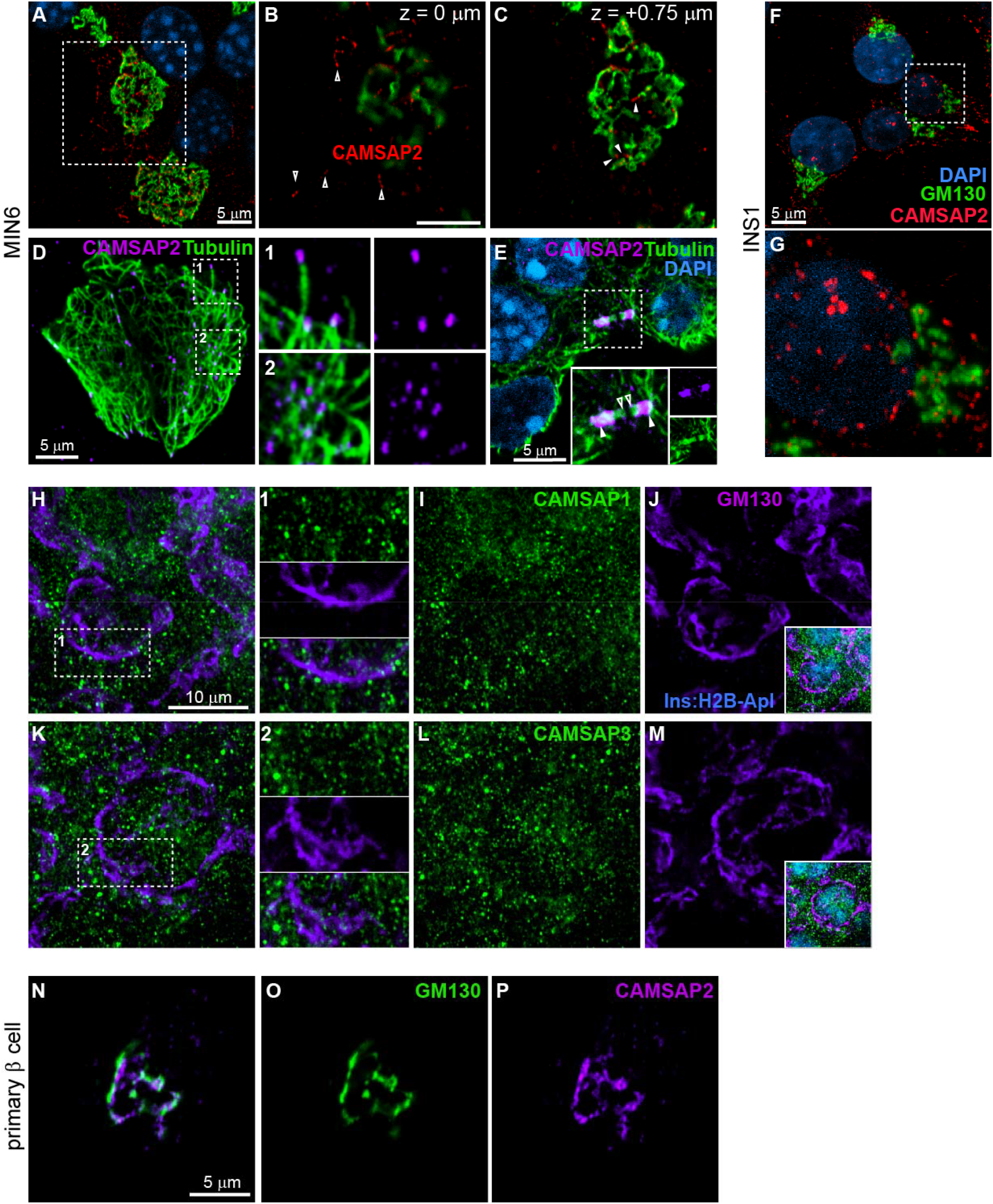
CAMSAP2 marks microtubule minus ends in MIN6 and in INS1 cells (A-C) Immunofluorescence staining of GM130 (green) and CAMSAP2 (magenta) in MIN6 cells. Blue, DAPI. (B) and (C) are at the same x-y position and 0.75 μm apart on the z-axis. (D-E) Immunofluorescence staining of tubulin (green) and CAMSAP2 (magenta) in MIN6. Blue, DAPI. Closed arrow heads, the minus ends of microtubules in the midbody; Open arrow heads, the plus ends. (F-G) Immunofluorescence staining of GM130 (green) and CAMSAP2 (red) in INS1 cells. Blue, DAPI. (H-M) Immunofluorescence staining of CAMSAP1 (green in H-J), CAMSAP3 (green in K-M) and GM130 (magenta) in mouse islet. Cyan, *Ins2* promoter-driven H2B-mApple. (N-P) Immunofluorescence staining of GM130 (green) and CAMSAP2 (magenta) in cultured primary β-cells dissociated from mouse islets.

**Figure S2.**
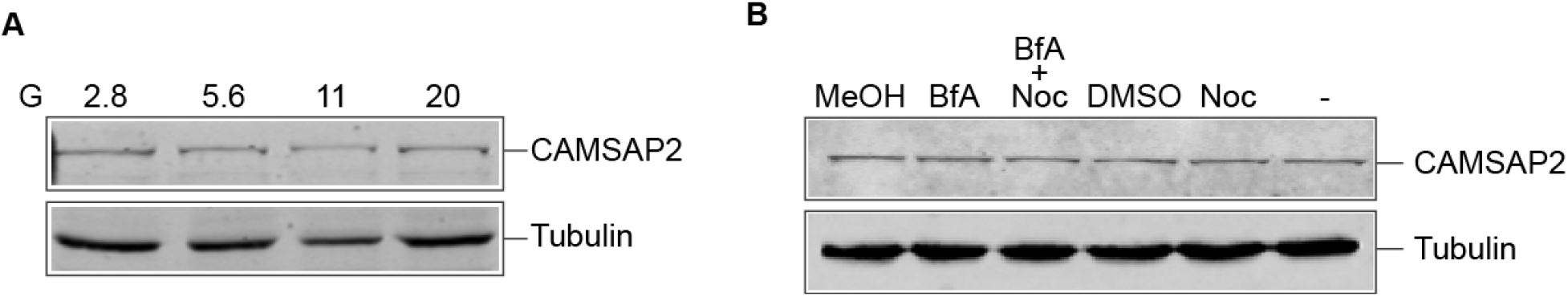
CAMSAP2 protein level is not changed by glucose stimulation or when the Golgi is collapsed. (A) Immunoblotting against CAMSAP2 and tubulin in lysates from mouse islets. Islets were incubated in media containing 2.8, 5.6, 11, or 20 mM glucose for 2 hours prior to lysis. (B) Immunoblotting against CAMSAP2 and tubulin in MIN6 cultured in media containing 25 mM glucose for 2 hours prior to lysis. Reagent includes 0.1% methanol or DMSO, 2.5 μM Nocodazole, and 5 μg/ml Brefeldin A.

**Figure S3.**
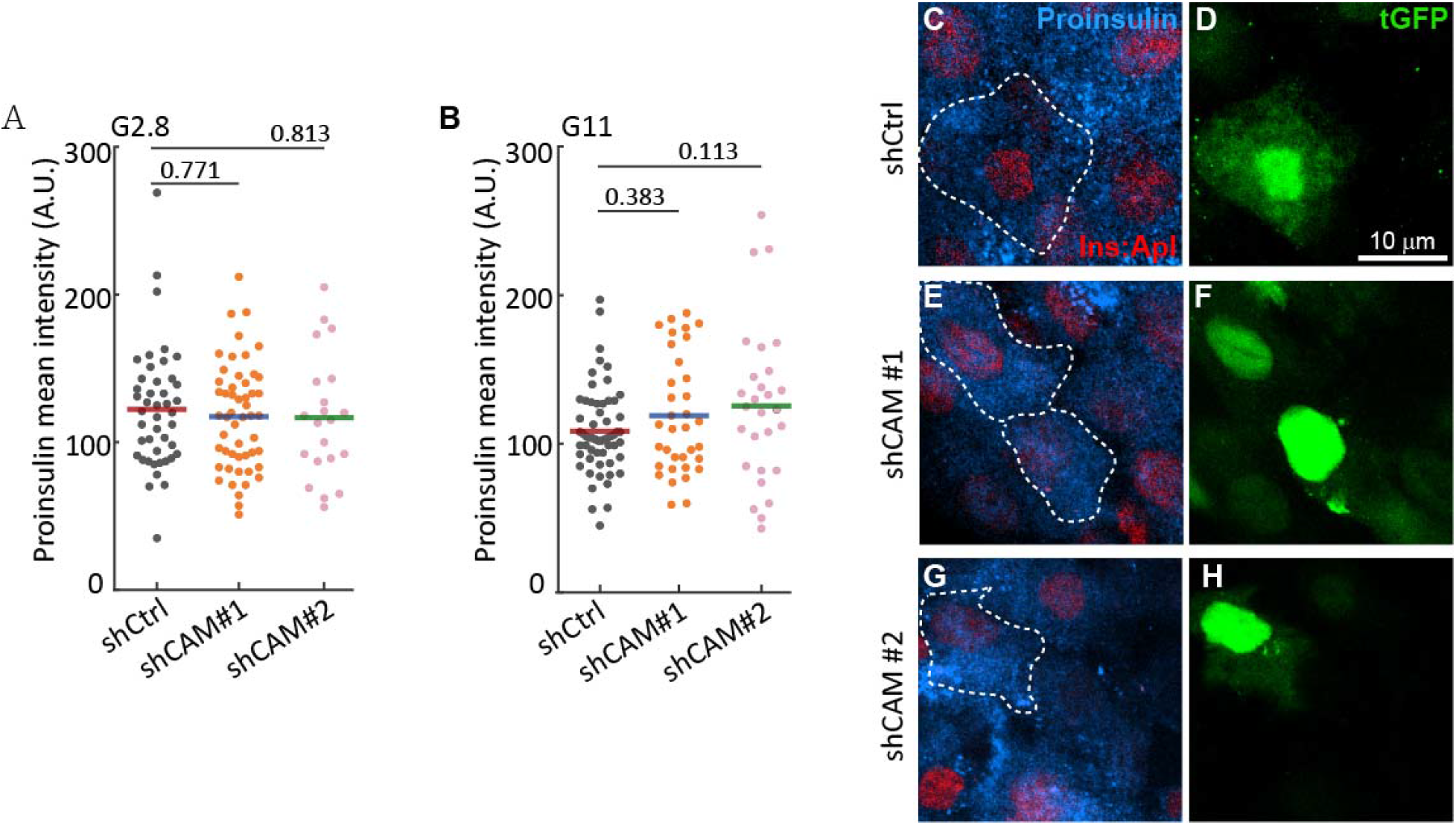
CAMSAP2 depletion does not alter the level of proinsulin in islet β-cells (A-B) Quantification of immunofluorescence intensity of proinsulin in β-cells in mouse islets cultured in media containing 2.8 mM (A) or 11 mM (B) glucose. Dots represent the mean intensity of proinsulin of individual cells. Bars represent averages from three animals. (Dunnett’s multiple comparisons test). (C-H) Immunofluorescence staining of proinsulin (cyan) in islets transduced with shRNAs and cultured in media containing 11 mM glucose. Dashed lines delineate transduced cells. Red, *Ins2* promoter-driven H2B-mApple; Green, tGFP to indicate successful transduction.

**Figure S4.**
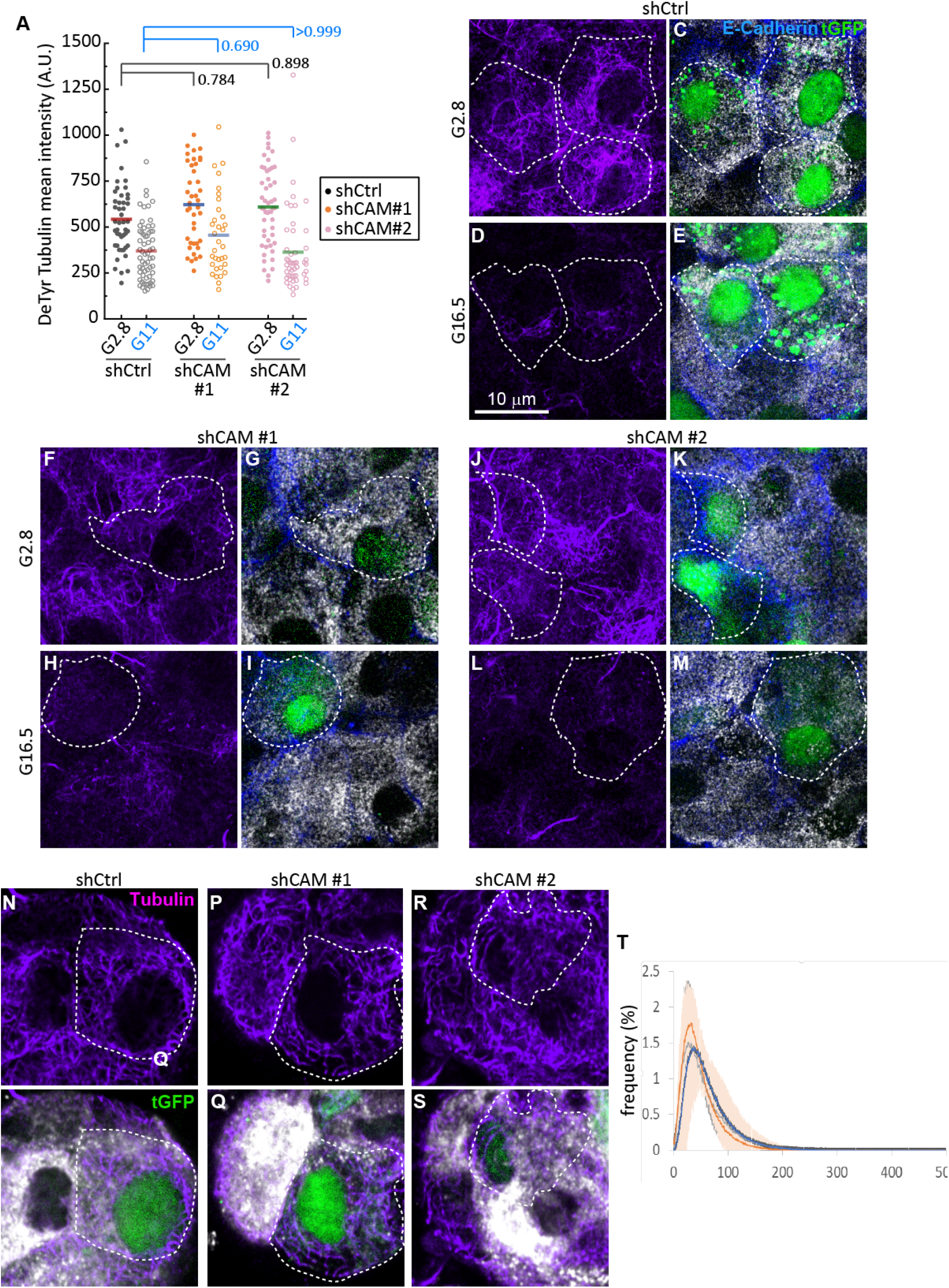
CAMSAP2 depletion does not interfere with microtubule dynamics in β-cells. (A) Quantification of detyrosinated tubulin intensity in transduced β-cells in mouse islets. Dots represent mean fluorescence intensity of individual β-cells in mouse islets. Bars represent the average of three animals. * p<0.05, *** p<0.001 (Dunnett’s multiple comparisons test). (B-M) Immunofluorescence staining of detyrosinated tubulin (magenta), E-Cadherin (blue), and insulin (white) in β-cells in mouse islets. Green, tGFP to indicate successful transduction. Dashed lines delineate transduced β-cells. (N-S) Immunofluorescence staining of tubulin (magenta) and insulin (white) in β-cells in mouse islets. Green, tGFP to indicate successful transduction. Dashed lines delineate transduced β-cells. (T) Quantification of tubulin distribution in the cytoplasm in transduced β-cells in mouse islets.

